# Universal high-throughput image quantification of subcellular structure dynamics and spatial distributions within cells

**DOI:** 10.1101/2024.08.18.608451

**Authors:** Andrew Martin, Sue Zhang, Amanda Williamson, Brett Tingley, Mira Pickus, David Zurakowski, Hadi T. Nia, Orian Shirihai, Xue Han, Mark W. Grinstaff

## Abstract

Image analysis of subcellular structures and biological processes relies on specific, context-dependent pipelines, which are labor-intensive, constrained by the intricacies of the specific biological system, and inaccessible to broader applications. Here we introduce the application of dispersion indices, a statistical tool traditionally employed by economists, to analyze the spatial distribution and heterogeneity of subcellular structures. This computationally efficient high-throughput approach, termed GRID (Generalized Readout of Image Dispersion), is highly generalizable, compatible with open-source image analysis software, and adaptable to diverse biological scenarios. GRID readily quantifies diverse structures and processes to include autophagic puncta, mitochondrial clustering, and microtubule dynamics. Further, GRID is versatile, applicable to both 2D cell cultures and 3D multicellular organoids, and suitable for high-throughput screening and performance metric measurements, such as half-maximal effective concentration (EC50) values. The approach enables mechanistic analysis of critical subcellular structure processes of relevance for diseases ranging from metabolic and neuronal diseases to cancer as well as a first-pass screening method for identifying biologically active agents for drug discovery.

**Statement of Significance:** Current methods for image analysis in microscopy are tailored to specific biological contexts, which creates challenges in implementation and efficiency for researchers studying a diverse range of subcellular processes. Our application of dispersion indices, traditionally used in economics, offers a universal framework for high throughput quantification of biological structures, enabling easier and more consistent analysis across various biological contexts. By transforming pixel intensity and count into meaningful statistical measures, our method simplifies the quantification of subcellular structures such as autophagic puncta, microtubule dynamics, and mitochondrial clustering. This approach accelerates the quantitative analysis of sub-cellular processes in disease as well as imaged-based drug discovery.

**Classification:** Bioengineering, Cell Biology, Applied Biological Sciences

## Introduction

Quantitative fluorescence microscopy is an indispensable technique for characterization of biological structures^1–4^, drug discovery and screening pipelines^5–7^, mechanism of action studies^8–10^, and studying disease pathology in preclinical studies^11–13^ to guide therapeutic development. While several methods and acquisition guidelines^14^ exist for image analysis of particular cellular organelles or structures, they rely on the specific biological contexts, which requires using multiple context-specific image analysis pipelines in ImageJ or in the subsequently improved software, CellProfiler^15^, for studying diverse biological features. We hypothesized that dispersion indices — mathematical measurements used to assess wealth distribution within a population — provide a universal framework for quantifying distributions without the need for tailored adaptations to distinct biological systems.

Dispersion indices are statistical measures of distribution regularly used in economics to measure income inequality within countries and across continents ^16,17^. Recognizing dispersion indices measure population disparities as a function of income and households gives rise to a new opportunity in quantifying biological structures in fluorescent microscopy images whereby pixel intensity and number are a proxy for income and households, respectively. Herein, we describe the use of dispersion indices to quantify the diffusiveness or aggregation of subcellular structures in images. The agnosticism of dispersion indices is advantageous to situations where a single quantification method must be applied to diverse biological contexts or when quantifying structures that lack established analysis protocols. We discuss the generalized entropy indices, GE(0) and GE(1), most commonly known as Theil’s L and Theil’s T, as well as the coefficient of variation (COV) and Gini coefficient. These indices have previously been used in biological contexts; the Gini coefficient, for example, has been used in biological contexts to analyze gene expression data^18,19^ and quantify bacterial aggregation^20^. We highlight the advantages of using dispersion indices to analyze the distribution of subcellular structures in biological images and demonstrate their adaptability across various experimental setups, in quantifying autophagic puncta, mitochondrial clustering, and microtubule polymerization. Dispersion indices effectively capture the nuances of subcellular structure distributions in diverse biological contexts, including an example in a multicellular 3D human midbrain culture.

Given the current challenges in quantifying autophagy^21^ and the clinical importance of autophagy in diseases^22^ such as diabetes, fatty liver disease, kidney disease, Parkinson’s, Alzheimer’s, and cancer, it is a prime test case to evaluate this new technique. Autophagic puncta are typically quantified via the protein marker LC3-II, which strongly associates to the membranes of autophagosomes and appears as puncta when stained. Despite LC3-II being spatially unique, the most common analysis method is via western blots, which lacks single cell resolution. This complicates the interpretations of autophagy in cell cultures and tissues that contain multiple cell types. Fluorescence-based methods offer single cell resolution and are commonly used in high content image analysis to segment cells using computer algorithms. However, the manual counting of LC3 puncta within images remains the standard method for quantitative analysis in autophagy research, as automated segmentation of autophagosomes^21,23^ from microscopy images is challenging. These challenges are highlighted in autophagy assay guidelines set by Klionsky et al.^24^, and a straightforward quantitative solution is still lacking. We present the development of a facile imaging-based assay as a platform for evaluating autophagy-modulating compounds and materials. Specifically, we describe the quantification of autophagic puncta using dispersion indices as an alternative to puncta counting. To demonstrate the universality of the dispersion indices approach, we then study mitochondrial clustering and microtubule dynamics, both of which are also of widespread interest.

## Results

### Dispersion indices simulation/validation

We adapted dispersion indices from their traditional use in economics to analyze the distribution of pixel intensities in images. By using dispersion indices in conjunction with image segmentation software, this analysis allows the quantification of intensity inequality within individual cells. In this application of dispersion indices, we substitute income for pixel intensity and households for pixels. Transfer of intensity from bright pixels to dim pixels decreases inequality measurements. The mathematical properties of dispersion indices are population and mean invariant – an attractive feature for image analysis. Regardless of the array size of the image (number of pixels) (Figure 1.A-D) and regardless of the mean of the distribution, images/objects which exhibit the same intensity distribution will yield the same value when using the same dispersion index. Thus, dispersion indices are intuitive and quantitative measures such that the indices increase when the distributed fraction decreases, meaning that the total pixel intensity distributes among fewer pixels (Figure 1.A-D). The analogy in terms of economic inequality means that the total income is distributed amongst a smaller number of people in the population, indicating higher inequality. However, it is important to note that it is necessary for the resolution of the image to be substantial enough to be able to expand the range of sensitivity to scenarios in which a high amount of the pixel intensity is distribution among a small fraction of pixels (Figure S1), as there are noticeable bounds on the maximum response of arrays which are smaller than 50 pixels. Consequently, the dispersion index is independent of cell size or resolution, as long as the cell is larger than 50 pixels. For example, an array of 2 pixels can only have a max inequality where 99.999% of the intensity is in 1 of 2 pixels, whereas an array of 50 pixels has max inequality when 99.999% of the intensity is in 1/50 pixels, although these cases are extreme. Additionally, dispersion indices anonymize pixels, such that the exact spatial distribution of the biological structure does not affect the dispersion index so long as the intensity distribution is unaffected.

**Figure 1.**
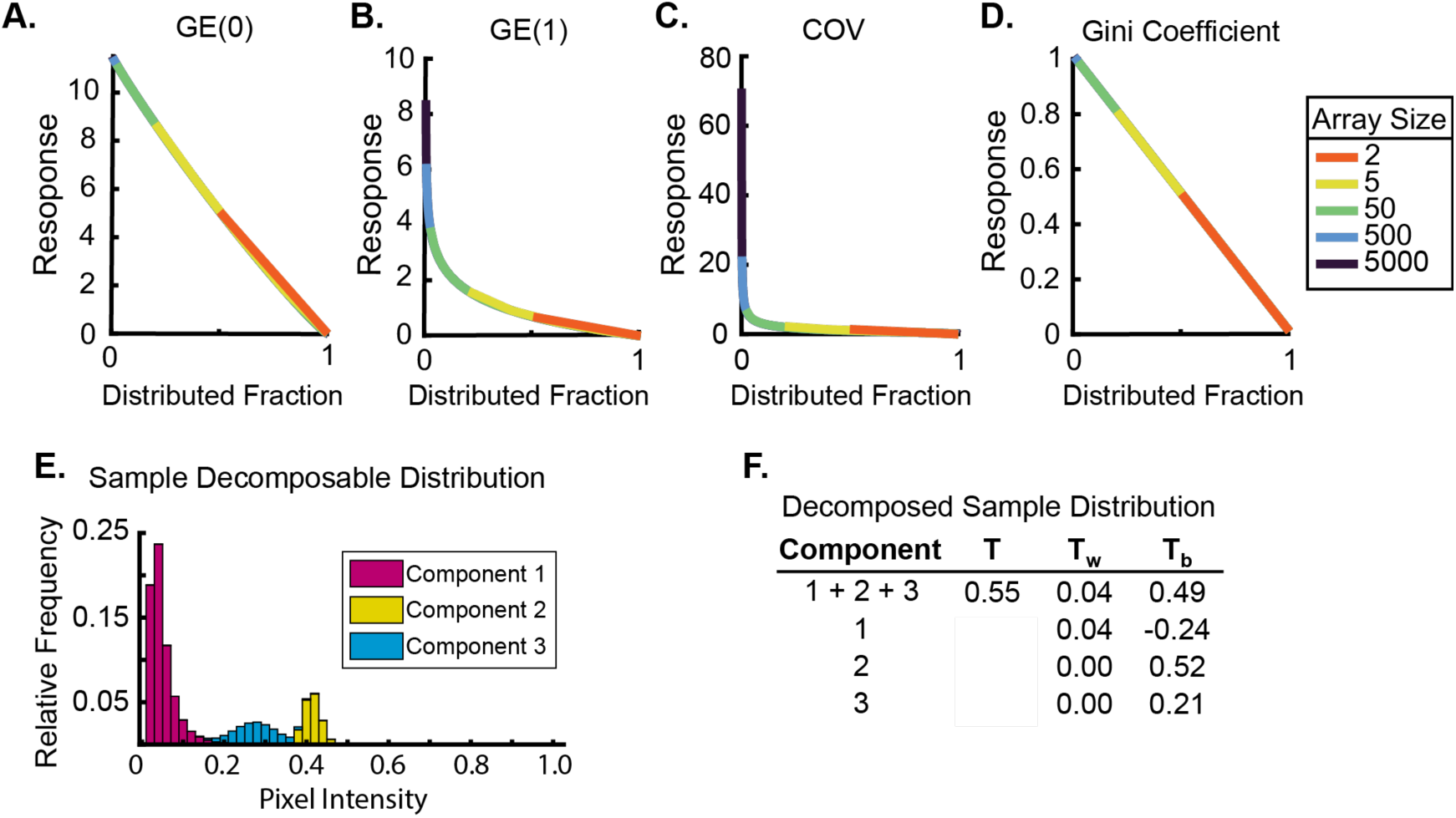
Numerical properties and decomposition of dispersion indices. (A-D) Measures of pixel intensity inequality using GE(0), GE(1), COV and Gini coefficient as a function of both i) the distributed fraction, which is the fraction of the pixel array that contains 99.999% of the total intensity, and ii) the array size, which is the number of pixels. For some plots, the array sizes are not plotted to facilitate ease of visualizing the trend. The plots converge as the array size increases and dispersion index responses follow similar trends irrespective of array size. E) Example histogram with an artificially-generated distribution of pixel intensities that can be linearly decomposed into three sub-components. The whole distribution is comprised of three sub-populations with different means, with two sub-populations having normal distributions. (F) Decomposition of this distribution using Thiel’s’ T index into sub-components reveals the Tw and Tb contributions from these features quantitatively.

Dispersion indices are uniquely sensitive to different fluctuations in inequality (Figure 1.A-D)^25^ . As noted within literature^26,27^, GE(1) (Figure 1.B) shows increased sensitivity over GE(0) (Figure 1.A) to changes among low populations of high income, which translates to bright pixels in image analysis indicating that the total intensity of the image is concentrated to a few pixels. Further, the curvature of the GE(1) plot shows a sharp increase in response as the distributed fraction decreases. Small changes at lower distributed fractions (i.e., the fraction of the pixel array that contains 99.999% of the total intensity, or the fraction of the population that contains 99.999% of the total income) result in large increases in the index response to those changes. The COV portrays the largest initial response to changes in these bright pixels. The Gini coefficient and GE(0) respond linearly to incremental redistribution of intensity. For GE(0) and Gini coefficient, there is an upper limit response value for max inequality, where one pixel contains 99.999% of the intensity (one bright pixel amongst dark pixels). However, for GE(1) and COV, the upper limit for measuring max inequality theoretically increases continuously as the array size increases. Additionally, generalized indices GE(0) and GE(1) decompose a distribution linearly into sub-components, such that sub-component contributions to overall index measures scale with the relative size of that sub-component (Figure 1.E,F)^16,17^. For example, a form of decomposition in measuring economic inequality would be separating the income distributions by age group to determine how each age group contributes to the overall population income inequality. This type of analysis is useful for images with multiple cell types. While Theil’s L and Thiel’s T are decomposable, it is not as intuitive to understand as the Gini coefficient, which is based on the Lorenz curve^28^.

### Simulation of dispersion indices within biologically relevant contexts

Autophagy is a natural protein degradation and clearance process that occurs at varying basal levels in mammalian cells. Much of autophagy research relies on the quantification of the protein marker, LC3-II, for autophagosomes^29^. During the formation process of autophagosomes (Figure 2.A), the phagophore sequesters cargo within a double membraned autophagosome and ultimately, the autophagosome merges with a lysosome to form an autolysosome. Enzymatic and acidic degradation of the inner membrane contents follows for subsequent recycling and use by cells. Wortmannin and bafilomycin A1 interrupt the natural autophagic process. Conversion of the cytosolic, unlipidated LC3-I to lipidated LC3-II results in the strong association of LC3-II to membranes of autophagosomes^29^, normally ranging 0.5 µm to 1.5 µm in diameter. LC3 lipidation affords increased puncta staining as LC3 is integrated into the autophagosomal membrane (Figure 2.B).

**Figure 2.**
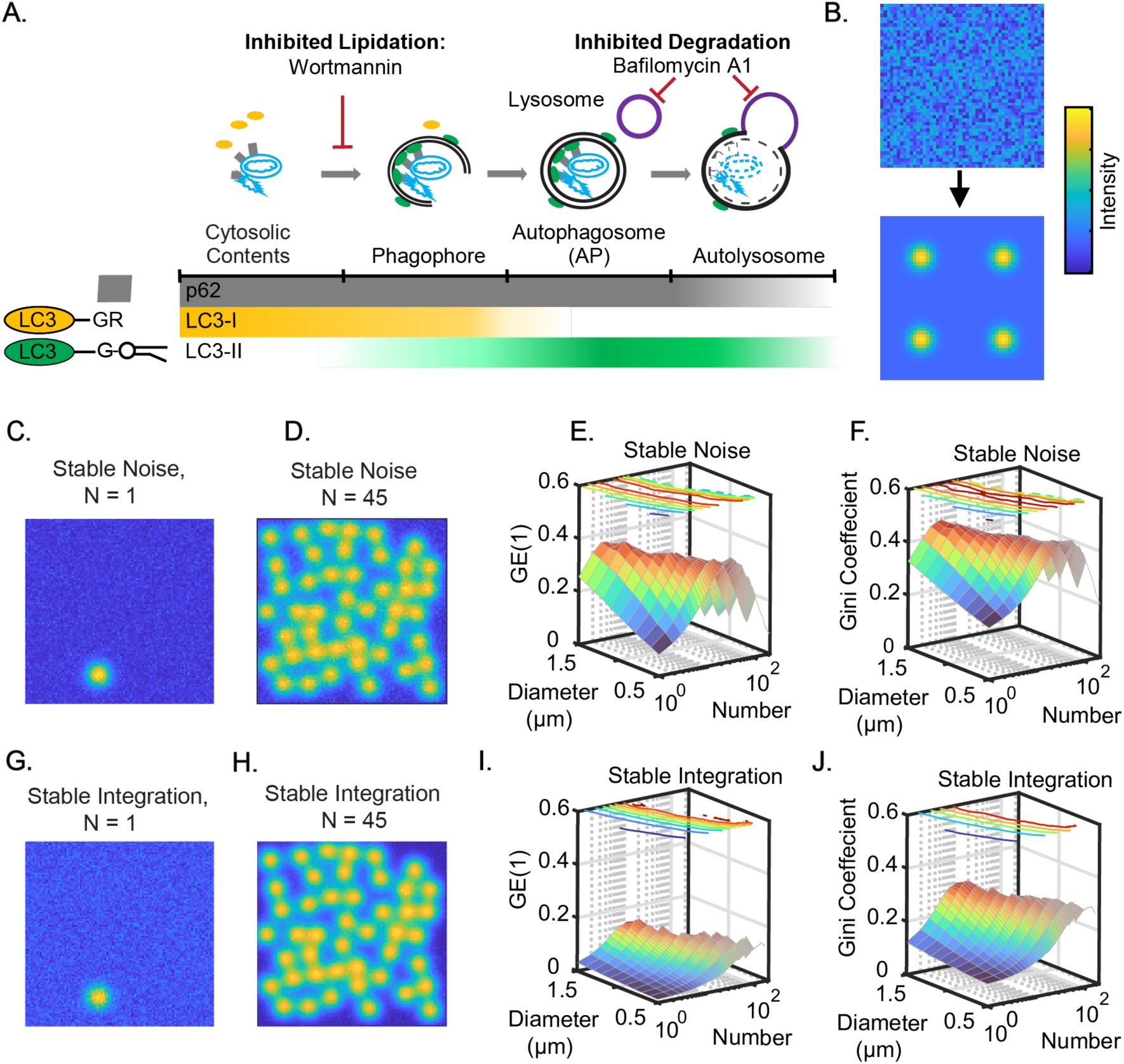
Numerical investigation into dispersion indices. (A) Cytosolic debris and proteins are sequestered into a double membraned autophagosome before degradation. LC3 and p62 mediate the formation process and are degraded. (B) Lipidation of LC3-I into LC3-II results in the spatial organization of LC3-II that results in bright puncta in images. (C,D) Example simulated puncta accumulation given a constant background signal intensity. (C) Input array when number of puncta = 1 and (D) at the maximum when the number of puncta = 45. (E,F) example 3D surface plots showing (E) GE(1) response and (F) is the Gini coefficient response from input arrays with varying puncta diameter and number of puncta. (G-J) represent the same simulations as (C-F) but under the assumption of stable total integrated intensity.

We investigated the statistical dispersion of the population in silico as a function of both increasing puncta number within the cell and puncta size. We considered two different simulation scenarios at biological extremes: 1) no loss of diffuse LC3-I background signal as [LC3-1] replenishes at an equivalent rate to conversion of LC3-II and the total LC3 content increases (stable noise) (Figure 2.C-F), and 2) diffuse LC3-I signal diminishes and total LC3 content remains constant (stable integration) (Figure 2.G-J). Simulation results reveal that high replacement of LC3 content (stable noise) yields a sharp and robust dispersion index responses and then peaks before decaying (Figure 2.E,F). The stable integrated intensity from constant LC3 levels affords increasing responses with puncta number and minimal effect due to puncta diameter, owing to the spatial focusing of pixel intensity to limited locations within the cell field (Figure 2.I,J).

Simulations of autophagosome accumulation in cells support the use of dispersion indices as a measure of puncta accumulation within cells. The two scenarios evaluated represent the theoretical bounds of LC3-I production rates, either allowing immediate replacement of LC3-I resulting in a stable background signal with an accumulation of overall intensity or limiting the replacement of LC3-I such that the overall intensity remains constant. In situations of no replacement and stable image mean, dispersion indices demonstrate nearly exclusive sensitivity to accumulating number of puncta rather than increasing puncta diameter, a finding that supports a high fidelity measurement. Furthermore, the simulation with high rates of LC3 replacement (stable noise) provides robust responses with the accumulation of low numbers of puncta, but with an eventual inflection point. We attribute this inflection point to the large changes in image properties such as image mean, a feature not represented in dispersion indices by design. Such an inflection point would be undesirable, but our simulation allows for the accumulation of puncta to the greatest extent which are non-overlapping, resulting in autophagosome numbers that are not found within live cells in a biological context. Additionally, we note that upon cell starvation, the lipidation of LC3-I is likely not equivalent to LC3-I replacement in contexts of high turnover, as simulated in the stable noise scenario. LC3-II is detectable via western blot within minutes of cell starvation^29^, but replacement of LC3-I depends on conversion from translated proLC3 and other post-translationally modified forms of LC3. As an example, non-autophagic, acetylated LC3B localizes to the nucleus and serves as a reserve normally and is depleted during starvation^30^. Therefore, stable integration more realistically simulates autophagic puncta in real biological contexts (Figure 2G,H).

### Generation of a biological dataset and evaluation of cell-based measurements

Given the robust response of dispersion metrics to the formation of simulated autophagosomes, we treated HEK293A and HFL cells to induce 4 different levels of LC3 lipidation and imaged the cells using a combination of antibodies and fluorescent fusion protein constructs (LC3B-GFP) (Figure 3.A-D,E). In parallel, we extracted protein content for western blot (Figure 3.F) to evaluate LC3 lipidation and p62 accumulation. In doing so, we determined the range of autophagic degradation capacity. Levels of p62 increase in nutrient deprived cells treated with Earle’s Balanced Salt Solution (EBSS). The PI3K Inhibitor, Wortmannin, inhibits LC3-I to LC3-II conversion as expected and Bafilomycin A1 treatment results in a block of autophagic degradation after LC3 lipidation, affording marked high levels of LC3-II in western blots and numerous puncta per cell. Quantification of LC3 channel images yields robust and reliable dispersion index responses to modulations in LC3 lipidation states and the presence of autophagosomes within cells (Figure 3.G-J). To determine the accuracy of dispersion indices as a quantitative measure of autophagy in cells, we analyzed regression models of all calculated image measurements against western blot results. A strong correlation (R^2^= 0.89) exists between the dispersion indices, GE(1) and Gini coefficient, and the LC3-II western blot data, confirming that dispersion indices align well with western blot results. The slope of the regression line indicates the sensitivity of each dispersion index to changes in LC3-II, where the regression line for the normalized Gini coefficient response has a higher slope of 2.31 (Figure 3.J) compared to a slope of 2.13 for the normalized GE(1) response (Figure 3.H).

**Figure 3.**
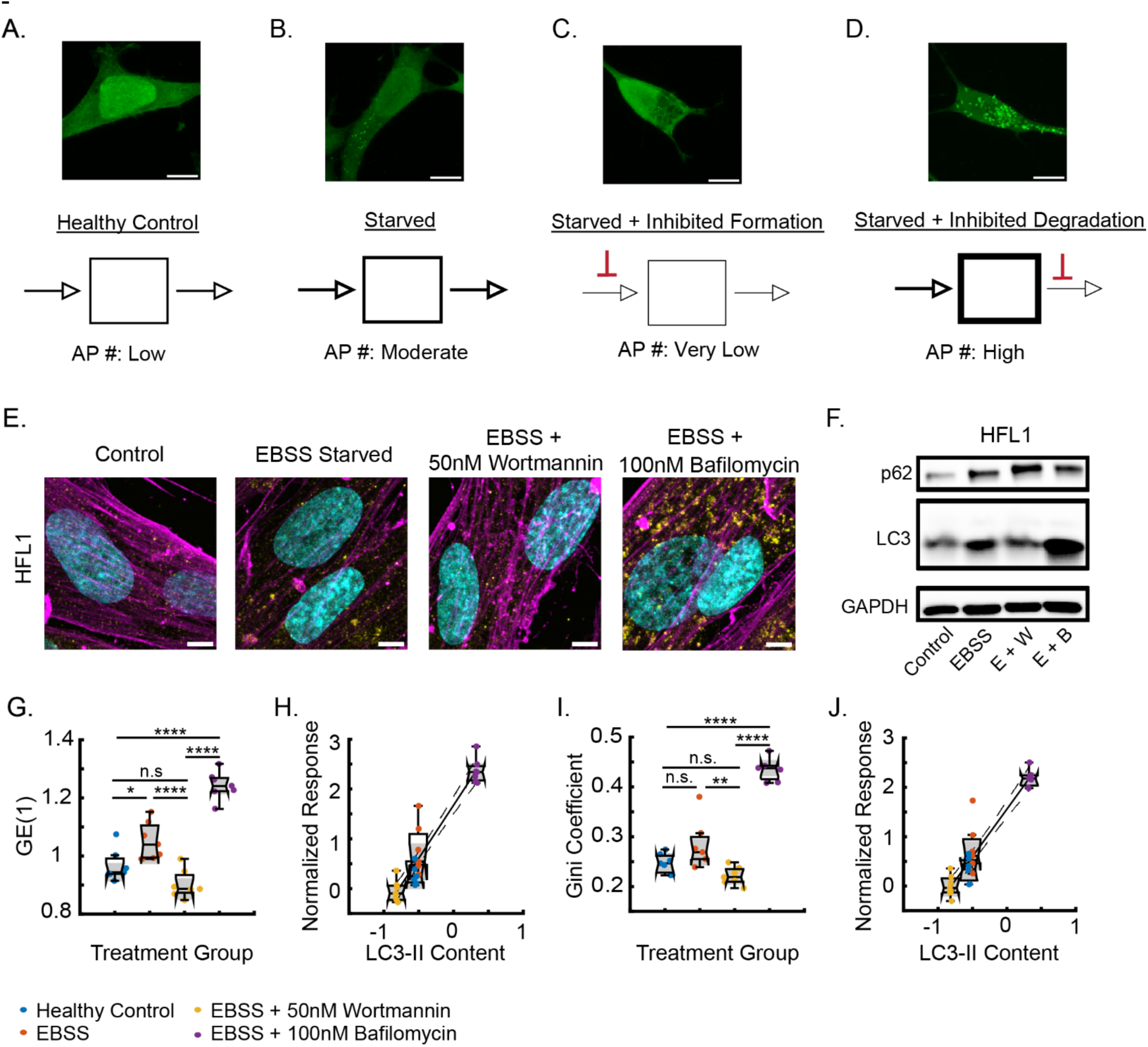
Dispersion Indices respond to altered lipidation states. (A) Heathy HEK293A cells expressing LC3-GFP have low autophagic flux and do not accumulate autophagosomes (AP). (B) Starved HEK293A cells increase the intracellular content recycling by increasing autophagic flux. (C) Starved HEK293A cells with inhibited LC3 lipidation via Wortmannin cannot form autophagosomes at the same rate, decreasing flux. (D) Starved HEK293A cells with inhibited degradation via 100nM Bafilomycin A1 cannot degrade autophagosomes. Autophagic flux is the difference between C and D under starvation. Scale bar is 5 µm. (E) Representative images of HFL1 cells under different states of autophagic flux. Cyan is Hoechst 33342; yellow is LC3 immunolabeling; magenta is β-actin. Scale bar 5 µm. (F) Representative HFL1 western blot data. (G) Plot of GE(1) treatment response for HFL1 cells. (H) Regression of normalized GE(1) against LC3-II western blot z-score results. (I) Plot of Gini coefficient treatment response for HFL1 cells. (J) Regression of normalized Gini coefficient against LC3-II western blot z-score results. Dotted lines represent the 95% confidence interval for the linear fit. * = p<0.05, ** = p<0.01, *** = p<0.001, **** = p<0.0001.

We generated a CellProfiler pipeline to segment and analyze autophagy images. We used CellProfiler to generate 285 different types of measurements per cell and to perform semi-automated LC3 puncta counting with manually selected imaging thresholds, as automated thresholding techniques were insufficient to reliably detect puncta in all treatment groups. We evaluated regression models for each of the measurements through 7 different characteristics (fit error, fit variability, fit sensitivity, treatment cross validation, KFold cross validation, R^2^ of fit, and adjusted R^2^) (Figure 4.A), and then evaluated the cell measurement types based on their performance scores. We removed models with insignificant fits before ranking to yield 40 different image measurements with significant correlations and predictive capability (Figure 4.B,C). Both cumulative and cell-type specific performance scores demonstrate that GE(1), GE(0), Gini coefficient, and COV are the best cell measurements investigated (Figure 4.C). To further visualize errors associated with image-based detection of LC3-II, we utilized regression error characteristic curves^11^ (Figure 4.D), plotting the fraction of data points (i.e., accuracy) as a function of increasing deviation from the model prediction (i.e., error tolerance). Other image-based measurements performed well but lacked the capacity to represent ∼20% of data. Other top performing cell-based measurements include LC3 puncta counting, as expected, and image texture measurements including image granularity. However, these other imaged-based features are unreliable in some cell types tested (Figure 4.C).

**Figure 4.**
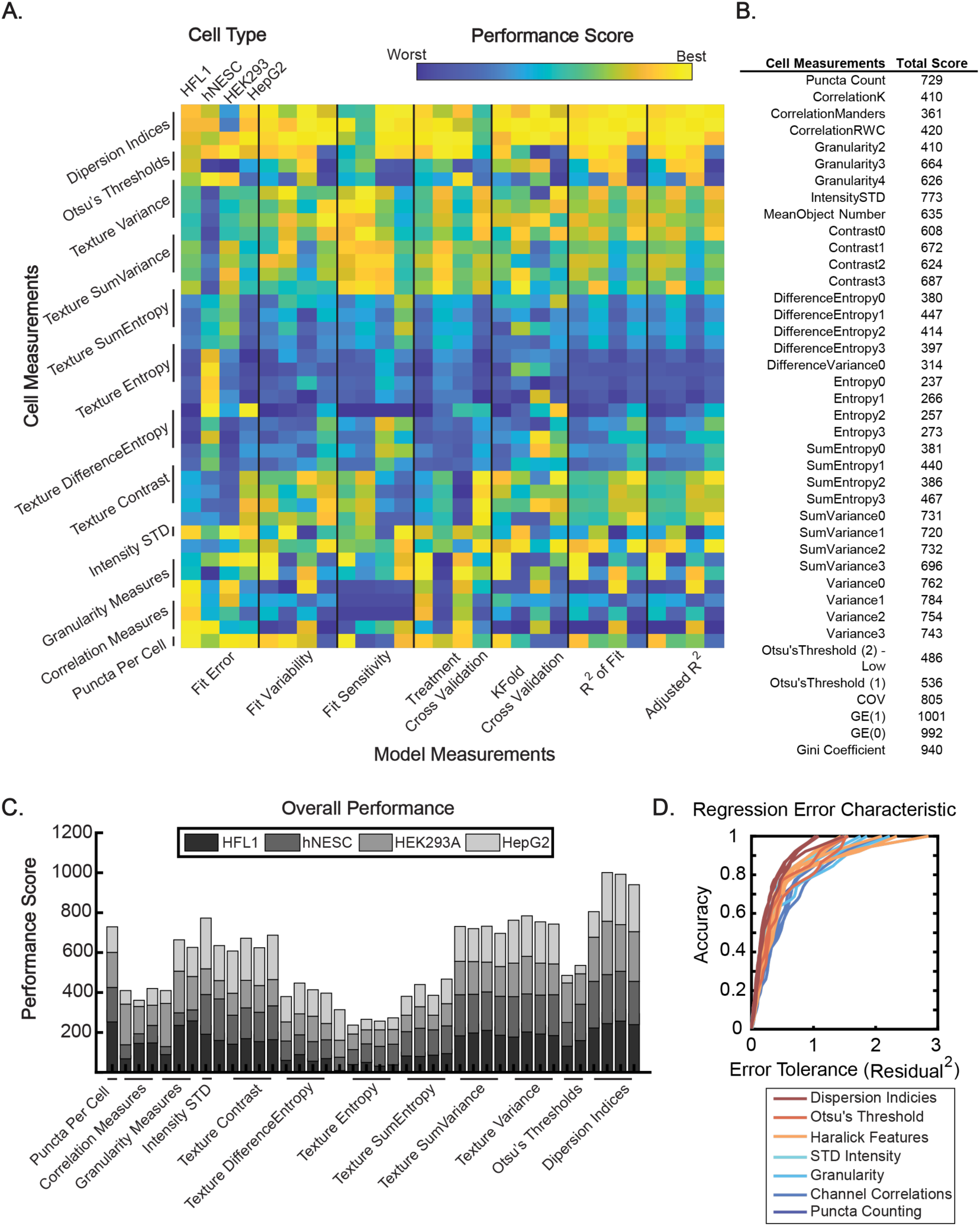
Rank-based performance model results. (A) Individual response scores for each cell measurement and model performance metric. (B) Table of overall top performing measurements. (C) Overall performance scores for evaluated metrics with significant correlations to LC3-II levels (p<0.05). (D) Regression Error Characteristic curves for parameters colored by calculation module.

### Decomposition of multicellular systems using GE(1) and GE(0)

Dispersion indices are highly useful parameters because they do not require object identification for any application – making them agnostic to the specific organelle or structure of interest. Moreover, generalized entropy indices contain the added benefit of linear decomposability, a feature shared with traditional puncta counting. In practice, the linear decomposability evaluates the effects that subgroups contribute to the overall inequality of the population, analogous to looking at the variability in puncta per cell measurements when averaged over a field of view. In the context of image analysis, decomposability determines if overall changes in the dispersion are due to differences in average intensity (e.g., caused by uneven staining) rather than changes in inherent dispersion within each of the subgroups.

To demonstrate the benefits of using the decomposability of dispersion indices GE(0) and GE(1) to identify inequality contributions by sub-components, we analyzed a multicellular human organoid culture model in response to starvation (Figure 5.A)^31^. Although derived from a single population of cells, human neuroepithelial stem cells undergo differentiation to yield multiple cell types over time, including neurons, astrocytes, and oligodendrocytes. (Figure 5.A,B). Starvation of organoids leads to a substantial increase in autophagic flux, as demonstrated by western blot analysis (Figure 5.C). LC3 flux increases in starvation conditions compared with bafilomycin A1 treated controls, but it is unclear which cells within the population are responsive. To investigate this, we analyzed two different cell types within the culture, neurons, identified by TUJ1 staining, and astrocytes, identified by GFAP staining.

**Figure 5.**
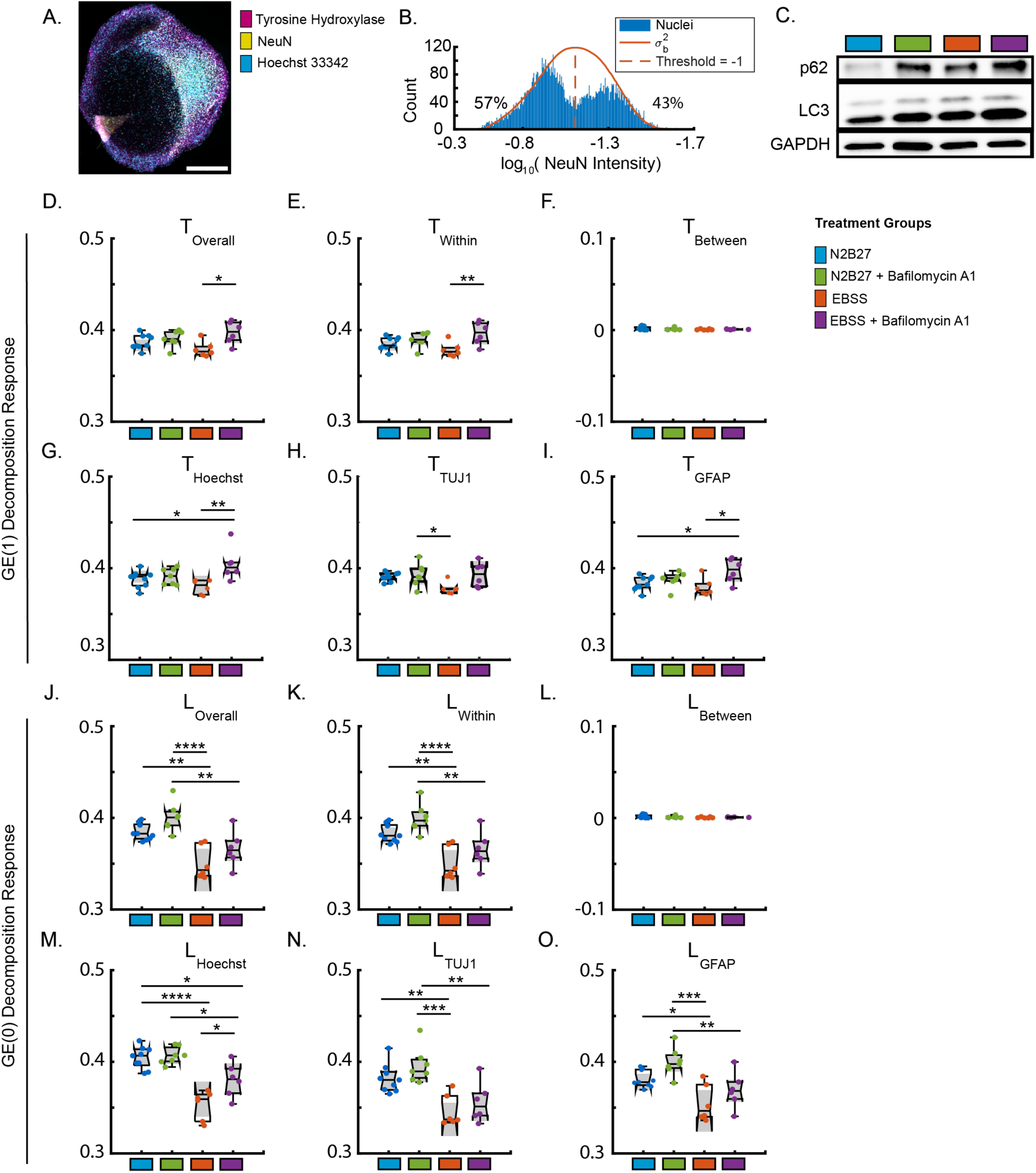
Decomposition of GE(1) and GE(0) in a multicellular system via cell-type specific markers. **(A)** 3D midbrain organoid slice labeled with tyrosine hydroxylase, NeuN and Hoechst 33342 to demonstrate multicellularity of the organoids. **(B)** Image quantification of slice in (A) as evidence for NeuN-and NeuN+ cell types within the 3D culture. **(C)** Western blot results from organoids under starvation conditions and inhibition of autophagosome degradation. **(D-F)** Decomposition of GE(1) to show within-group and between-group effects. **(G-I)** GE(1) T_within_ responses for nuclei, TUJ1+ neurons, and GFAP+ astrocytes, respectively. **(J-L)** Decomposition of GE(0) to show within-group and between-group effects. **(M-O)** GE(0) L_within_ responses for nuclei, TUJ1+ neurons, and GFAP+ astrocytes, respectively. * = p<0.05, ** = p<0.01, *** = p<0.001, **** = p<0.0001. All statistics are from ANOVA with multiple comparisons post hoc. Each point represents 1 FOV. N=3 organoids.

We first used the linear decomposition of GE(1), as it yields the best approximation of LC3-II levels within cells tested. Decomposition results of TUJ1 and GFAP stained cells reveal even LC3 staining between groups (Figure 5.F). The majority of the variation in staining intensity between treatment groups arises from differences within each cell type subgroup (Figure 5.E). For all stained areas, GE(1) reveals an increase in accumulated LC3 in starved cells (in EBSS) inhibited with bafilomycin A1, a response that follows western blot results (Figure 5.D). Interestingly, subgroup analysis using GE(1) shows a cell-type specific response to starvation, with TUJ1+ areas decreasing GE(1) response upon starvation, with bafilomycin treatment exhibiting little effect on the absolute magnitude of responses (Figure 5.H). This result shows that TUJ1+ areas had insignificant changes to autophagic flux due to starvation. In contrast, GFAP+ cells show elevated GE(1) response (i.e., LC3-II accumulation) in starved (EBSS) cells when treated with bafilomycin A1, revealing that astrocytes are starvation-responsive cells (Figure 5.I). The Hoechst subgroup yielded results similar to GFAP+ cells (Figure 5.G). These results align with literature showing that starvation conditions afford less pronounced effects in neurons than in astrocytes, where autophagy is robustly activated^32^. Finally, we evaluated these cells using the linear decomposition of GE(0), which enables more equal analysis of pixels with varying intensities. Similar to GE(1), the differences between groups minimally contributes to the overall differences between treatment groups. However, overall and cell-specific subgroup analyses show an increased GE(0) response between cells under fed conditions (Figure 5.J,M-O). The measured effects are most pronounced in Hoechst associated pixels, and similar for GFAP+ and TUJ1+ cell areas. In sum, Hoechst+, TUJ1+, and GFAP+ areas demonstrate a response to both starvation and to inhibition of autophagosome degradation, but in different ways.

Decomposition enables 3D autophagy measurements without the need for image-based object detection in multicellular systems. Indeed, dispersion indices reveal cell-type specific responses that are not represented through bulk measurements. Specifically, TUJ1+ neurons display a decreased level of dispersion upon starvation, potentially increasing autophagosome clearance in response to the stimulus but not accumulating LC3-II like neighboring GFAP+ cells. It is likely that basal autophagy is high in these neurons, as much of the differentiation factors included (e.g., BDNF, GDNF, cyclic AMP) can induce autophagy. The analysis of complex multicellular systems will greatly benefit from this type of analysis.

### Dispersion indices to quantify autophagosomal EC50

Next, we demonstrated the utility of dispersion indices for reliable, reproducible estimations of EC50 concentration within cells. In particular, we chose COV to measure EC50 due to the ease of access to the standard deviation and mean of the cell’s intensity through all image-processing software. We used a commercially available HEK293A GFP-LC3B line and cultured these cells with bafilomycin A1 under normal and starvation conditions to determine an EC50 value for autophagosome formation (Figure 6.A,B). We labeled nuclei and actin filaments to easily identify cells using high-resolution confocal microscopy (Figure 6.C,D). In this first group of experiments, we confirmed dose-response characteristics of these cells. We measured autophagy by two methods, the coefficient of variation (COV) for each cell as well as the number of LC3 puncta identified as objects. Our imaging results estimate an EC50 of ∼1nM for bafilomycin A1 in cells cultured under normal conditions and an EC50 of ∼4 nM for bafilomycin A1 in cells cultured under starvation conditions, which agrees with published results^33^. Moreover, the starvation of HEK293A cells raises the response ceiling relative to complete media – evidence for increased autophagic flux. This increase in autophagic flux is in tandem with increased cellular resistance to the inhibition of lysosomal degradation, with a shift of EC50 from ∼1.5 nM to ∼3.4 nM. This shift is also apparent upon visual inspection of the images (Figure 6.C,D), and most apparent is the shift of intracellular COV intensity, as opposed to puncta per cell measurements. Additionally, the use of widefield microscopy greatly reduces image acquisition time of the dataset. However, we observed that widefield microscopy increases the variability of the acquired dataset, where EC50 is reported as 7.6 ± 1.4 nM (95% confidence interval) when quantified using COV and the EC50 is reported as 5.1 ± 1 nM (95% confidence interval) when quantified using puncta counting (Figure S2). While our switch from confocal to widefield microscopy does not reduce our theoretical XY resolution, the loss of z-sectioning likely affects the variability of the widefield dataset.

**Figure 6.**
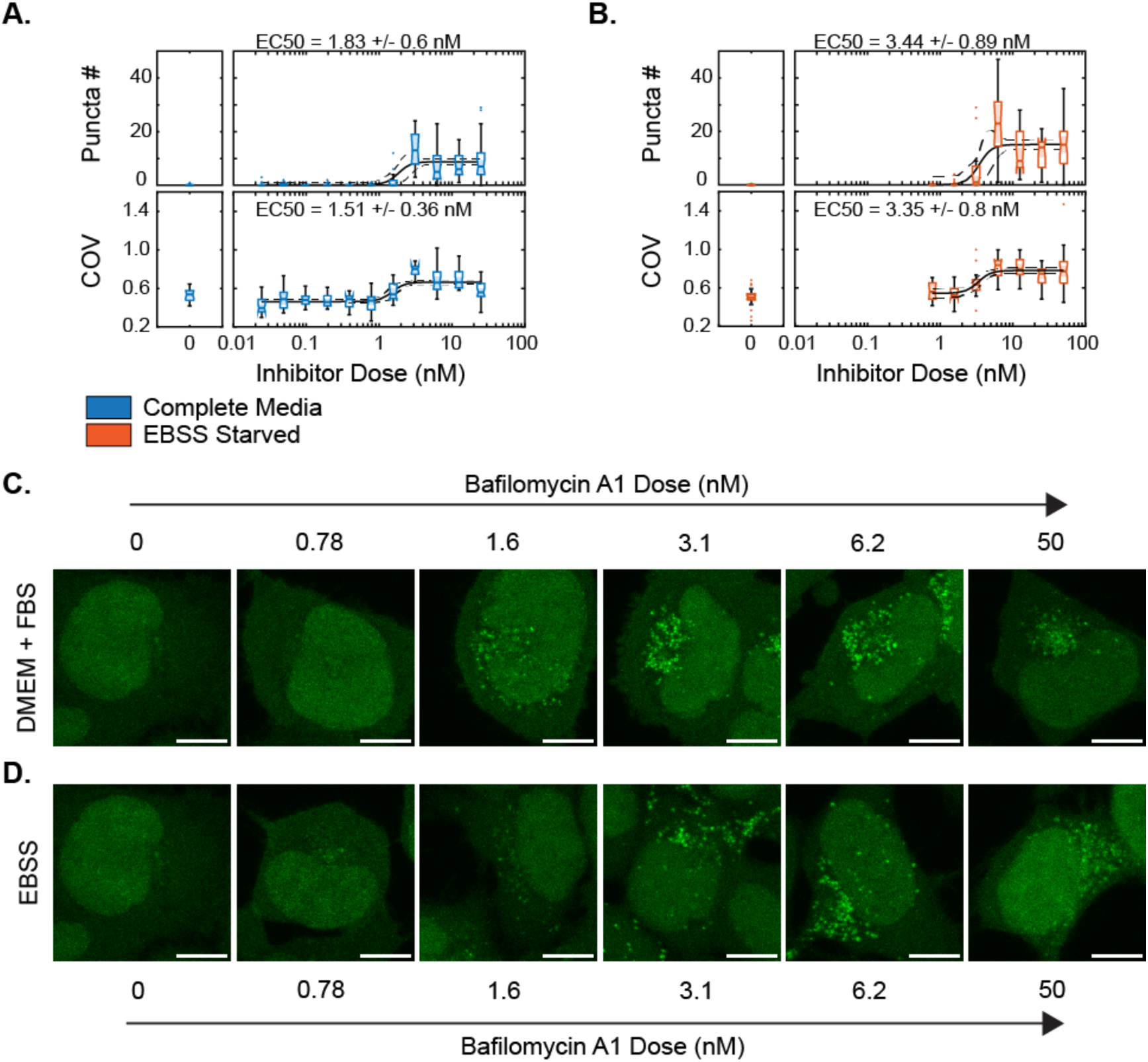
Starvation-dependent dose-response of HEK293A cells to bafilomycin A1. (A) Puncta per cell measurements (top) and coefficient of variation (COV) measurements (bottom) of HEK293A cells treated with bafilomycin A1 for 3 hours under complete media. (B) Puncta per cell and COV analysis for HEK293A cells under EBSS starvation for 3 hours with bafilomycin A1 treatment. Representative HEK293A GFP-LC3B images for cells treated with bafilomycin in (C) complete media and (D) in EBSS (starved cells). Images were acquired using confocal microscopy. Scale bar 5 micron. EC50 values are reported with a 95% CI.

### Dispersion indices to quantify mitochondrial clustering and microtubule dynamics

Next, we investigated applicability of dispersion indices to two other important biological structures: mitochondria and tubulin. Mitochondria are crucial for cellular energy production and are involved in various cellular pathways. Efficient microtubule dynamics are essential for cell division, and standard of care cancer treatments utilize microtubule inhibitors like paclitaxel as chemotherapies. Common practices for analyzing these structure require tedious processes such as mitochondrial isolation and western blotting. We hypothesized that quantitative assessments of mitochondria and tubulin via image analysis provide a facile pipeline for studying metabolic disruption and identify emerging cell resistance to chemotherapeutic drugs.

Literature reports that under hypoxic conditions, mitochondria cluster in the perinuclear region^34^ as quantified by measuring the fluorescence intensity of mitochondria at different distances from the nucleus (Figure 7B). However, dispersion indices enable the quantification of mitochondrial clustering without requiring detailed spatial information. All four dispersion indices show lower dispersity of mitochondria in hypoxic conditions, leading to higher dispersion index values (Figure 7C). Dispersion indices identify cells under normoxic and hypoxic conditions based on mitochondrial clustering.

**Figure 7.**
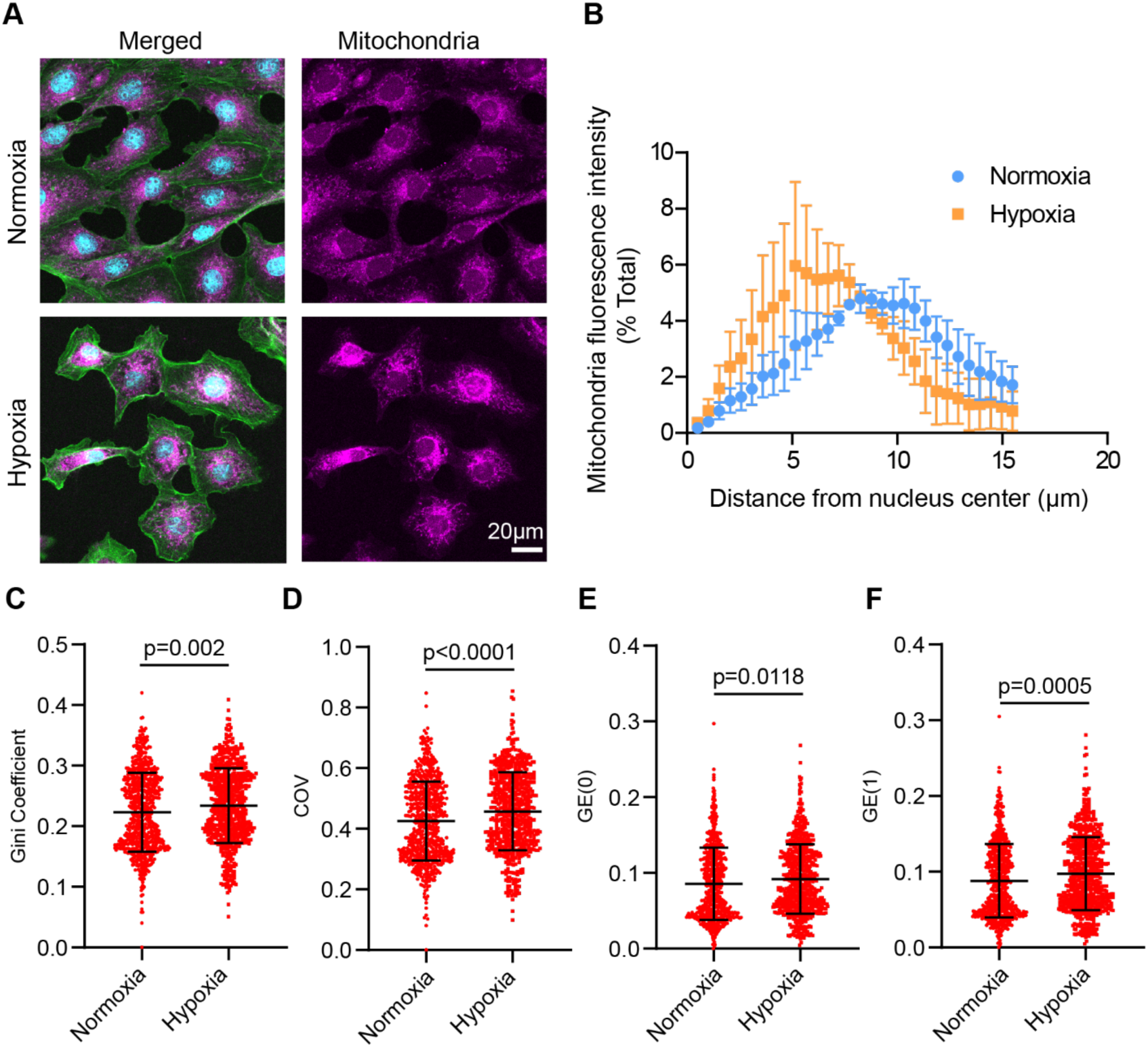
Quantification of mitochondrial clustering in normoxic and hypoxic conditions using dispersion indices. (A) Representative images of rat pulmonary artery cells stained for nuclei (cyan), actin (green), and mitochondria (magenta) in normoxic and hypoxic (2% O_2_) conditions. (B) Intensity profile of mitochondria as a function of distance from the center of the cell in normoxic and hypoxic conditions. (C-F) Quantification of mitochondria using Gini coefficient, COV, GE(0), and GE(1). Statistics are from unpaired two-tailed t-tests. N=659-664 cells.

Vincristine, an anti-mitotic microtubule destabilizing chemotherapy drug, binds to both soluble and microtubule-associated tubulin^35^. A previous study showed that vincristine alters the Haralick texture homogeneity^36^, which is an image analysis method used to analyze the relationship of neighboring pixels^37^. This finding prompted us to investigate if dispersion indices will quantify tubulin distribution. At concentrations of 100nM and 1μM, the dispersion index is significantly lower for all 4 dispersion indices. These results align with previous image analysis reports using Haralick texture homogeneity showing that 1μM significantly alters tubulin distribution, which leads to increased homogeneity. Tubulin is more evenly distributed, which is consistent with our finding that 1μM vincristine results in lower dispersion indices compared to the control. Interestingly, the Gini coefficient identifies cells treated with vincristine concentrations of 10μM and 20μM to be significantly different from the control, although the other three dispersion indices do not indicate significant differences. The observed discrepancy is likely attributed to a greater than 25% reduction in nuclei diameter, indicating reduction in cell size at 10μM and 20μM, resulting in loss of spatial distribution. The dispersion indices of tubulin treated with 10μM and 20μM vincristine become more similar to that of control cells. As the cell size decreases (Figure 8F), pixel intensities artificially appear more clustered and cause an artificial increase in the dispersion index. Additionally, higher concentrations of vincristine can result in formation of crystalline tubulin aggregates^38^ that appear punctated, contributing to the increase of the dispersion index. Dispersion indices offer advantages over Western blot, which examines soluble and polymerized forms of tubulin, but lacks the ability to assess time dynamics in a single cell. However, based on our findings, we note that the use of dispersion indices to compare treatments should be used with caution when there is a greater than 25% change in cell size due to treatment.

**Figure 8.**
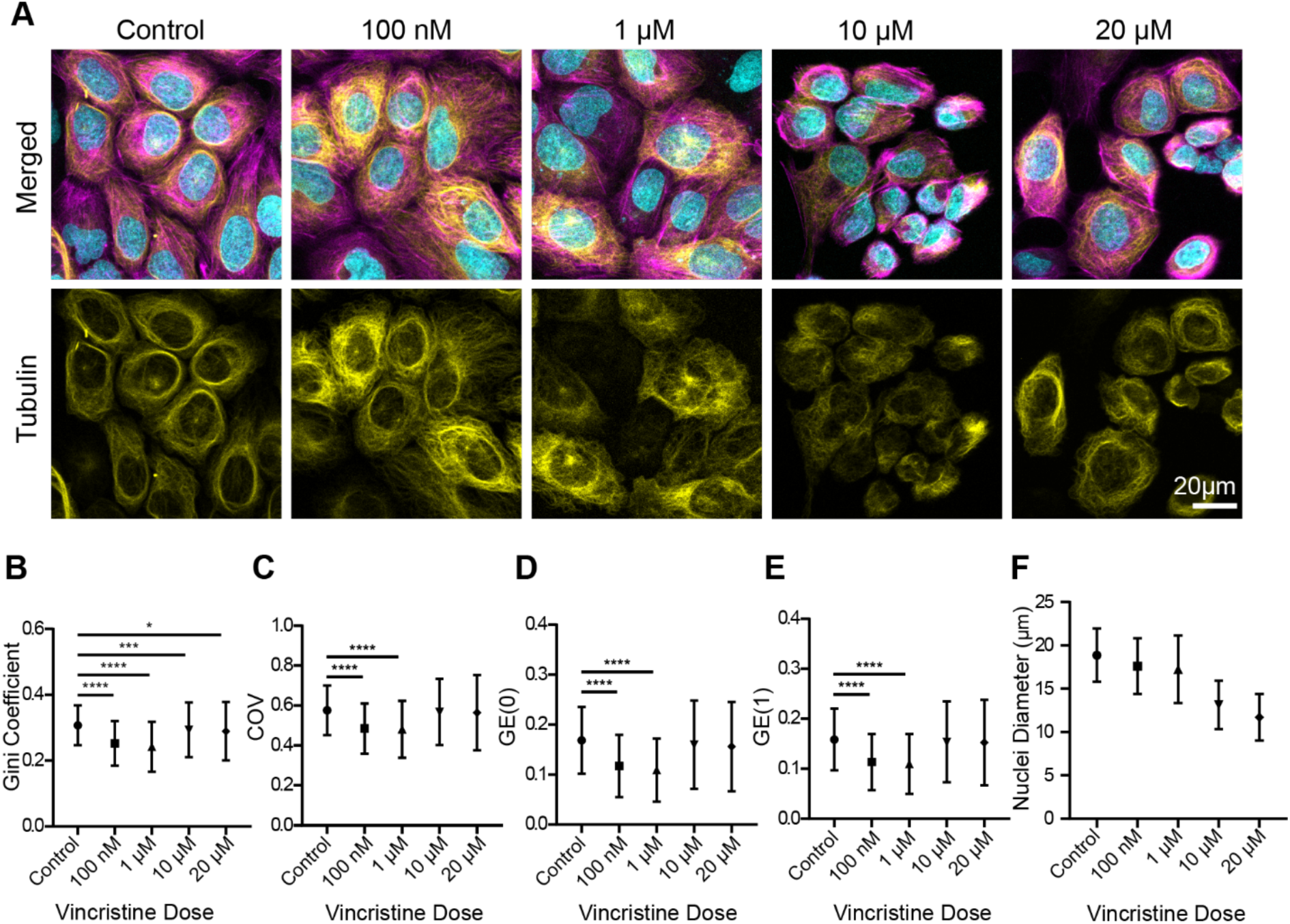
Quantification of tubulin dynamics with varying vincristine dosages using dispersion indices. (A) Representative images of U2OS cells stained for nuclei (cyan), actin (magenta), and tubulin (yellow) at concentrations of 100nM, 1μM, 10μM, and 20μM of vincristine for 30 minutes. (B-E) Quantification of tubulin using Gini coefficient, COV, GE(0), and GE(1). (F) Quantification of nuclei diameter. Statistics are from one-way ANOVA with Dunnett’s multiple comparisons test. N=137-864 cells.

## Discussion

We describe a novel image analysis paradigm that, while initially demonstrated using autophagy, is broadly applicable to other cellular structures. Taking inspiration derived from quantitative analyses of economic income inequality, we substitute pixel intensity for income and number of pixels for population and perform a comprehensive evaluation against hundreds of image measurements from open-source image analysis software. GE(1) (Theil’s T) provides the most effective model of LC3-II levels, followed by the GE(0), Gini Coefficient, and the Coefficient of Variation.

GE(1), which describes the entropic distance between the current observed entropy and the maximum possible entropy in the image array, measures the spatial focusing of LC3 to autophagosomal membranes, reflecting the reduced entropic states when LC3 is lipidated. Importantly, these quantification strategies outperform custom semi-automated puncta identification algorithms, despite the latter’s accuracy in representing LC3-II levels (Figure 4.B). Utilizing dispersion indices offers a significant advantage in terms of time saved compared to the tedious optimization required for object detection algorithms.

While image textural features provide another set of valuable measurements, the indirect nature of the best textural measurements (i.e., Variance and SumVariance) of autophagy as pixel intensity dispersion indicators underscores the utility of dispersion indices. Previous methods reliant on puncta identification, such as “autophagosome scores,” have been used to calculate a ratio between the total intensity within identified autophagosomes and the total intensity within the cell^39^. Dispersion indices are a similar type of measurement, but eliminate the need to identify individual puncta.

Applying this analysis paradigm to multicellular 3D human midbrain cultures, we observe differential cell responses between neurons and astrocytes upon nutrient deprivation. This is the first simultaneous measurement of autophagic flux in multiple cell types. We also demonstrate the applicability of our methods by generating EC50 curves for autophagosome degradation inhibition using Bafilomycin A1 under starvation. Our results underscore the method’s versatility and applicability across diverse biological contexts.

Moreover, our method extends beyond autophagy to analyze mitochondria and tubulin, which are vastly different structures from autophagosomes. This generalizability enables high-throughput screening of multiple biological structures, offering a powerful tool for current and future imaging assays. We anticipate our analysis pipeline to be an effective and versatile method in high content image analysis, where previously challenging targets can be quantified without the use of proprietary software or image analysis algorithms limited to specific cell types.

## Materials and Methods

### Cell Culture

HEK293A cells expressing LC3-GFP were purchased from Sigma-Aldrich (14050801). HepG2 (HB-8065), HFL1 (CCL-153), and U-2 OS cells were sourced from ATCC. Rat primary pulmonary artery endothelial cells (PAECs) were purchased from Cell Biologics (RA-6059). HEK293A, HepG2, and HFL1 cells were cultured with Dulbecco’s Modified Eagle’s Medium (Gibco, 11-965-092) supplemented with 1% penicillin-streptomycin and 10% Fetal Bovine Serum (Atlanta Biologicals, S11150). PAECs were cultured using the Complete Endothelial Cell Medium Kit (M1266, Cell Biologics). U-2 OS cells were cultured with McCoy’s 5A Medium (ATCC) supplemented with 1% penicillin-streptomycin and 10% Fetal Bovine Serum. Cells were grown at 37°C and 5% CO_2_. Cells were grown to ∼80% confluency and then split onto glass coverslips (for confocal microscopy), or to 96 well plates (CellVis) at a density of 4000 cells/well. Cells were left to grow on glass for a minimum of 48 hours before experiments began.

### Human Midbrain Organoid Culture

Human neuroepithelial stem cells (hNESCs) cells were a gift from Jens Schwarmborm at the University of Luxembourg and grown according to published procedures^31^. Stem cells were grown using N2B27 base media with, 150 μM ascorbic acid (Sigma-Aldrich, A59060), 0.75 μM Purmorphamine (Sigma-Aldrich, SML0868), and 3 μM CHIR-99021 (Axon Med Chem, 1386). hNESCs were grown on Matrigel-coated 6 well plates and media was changed every other day. To form organoids, hNESCs were seeded at 9000 cells/ well into a round-bottom 96-well plate (Corning, 7007). Single colonies were transferred to 24 well plates 4 days after seeding and embedded in a 30µL Matrigel droplet. Organoids were differentiated in N2B27 Media with BDNF (Peprotech, 450-02), GDNF (Peprotech, 4510-10), TGF-β3 (Peprotech, 100-36E), N^6^,2-O-Dibutyryladenosine 3’,5’-cyclic monophosphate sodium salt (D0627, Sigma-Aldrich), and ascorbic acid. Purmorphamine was kept in the differentiation medium for the first 6 days and then removed. Media was changed 3 times per week for 8 weeks before organoid use in experiments, and embedded organoids were kept freely floating in 24 well plates on an orbital shaker. N2B27-based media is a 1:1 (v:v) mixture of neurobasal medium (Gibco, 2110349), and DMEM/F12 Mixture without L-glutamine, (Corning, 15-090-CV), supplemented with GlutaMAX (Gibco, 35050061), N2 (Gibco, 17502001), and B27 (Gibco, 17504044).

### Nutrient Depravation and Autophagy Inhibition

Nutrient deprivation of cells and organoids to induce autophagosome formation involved the replacement of complete media with Earle’s Balanced Salt Solution (Gibco) for 3 hours with the addition of either DMSO (as mock control), Wortamannin (Sigma Aldrich) or Bafilomycin A1 (Medchem express). All inhibitors were added at the induction of starvation.

### Hypoxia Experiments

PAECs were seeded on 15 mm glass coverslips in 60mm petri dishes at a density of 0.8×10^6^ cells/dish for 48 hours before exposure to hypoxia. The petri dish was placed in a hypoxia chamber at 2% O_2_ for 2.5 hours and incubated at 37°C. The petri dish was then taken out to stain for mitochondria using MitoTracker Deep Red FM (Thermo Fisher Scientific) and placed back into the hypoxia chamber at 2% O_2_ for an additional 30 minutes. The cells were then stained for actin using CellMask Green Actin Tracking Stain (Thermo Fisher Scientific) and nuclei using Hoechst. Cells were fixed for 10 minutes at RT with freshly diluted 4% PFA in 1x PBS. Cells were rinsed and then mounted on glass slides using Prolong Diamond Antifade reagent (Thermo Fisher Scientific).

### Tubulin Inhibition Experiments

U-2 OS cells were seeded on glass-bottom 12 well plates at a density of 0.1×10^6^ cells/well for 48 hours before treatment. Cells were stained for tubulin using Tubulin Tracker Deep Red (Thermo Fisher Scientific), for actin using CellMask Green Actin Tracking Stain (Thermo Fisher Scientific) and nuclei using Hoechst. Cells were treated with vincristine (Sigma-Aldrich) solubilized in DMSO and FluoroBrite DMEM (Gibco).

### Autophagy Staining

For HEK293A cells, wells were fixed for 10 minutes at RT, with freshly diluted PFA to achieve a final concentration of 4% PFA in 1X PBS. Cells were then washed once with PBS, and then incubated for 30 minutes with Phalloidin iFluor 647 (AAT Bioquest, 1:10,000 dilution), Hoechst 33342 (1µg/mL), and 0.1% Triton-X 100 (Sigma Aldrich) in PBS. For cells grown on coverslips, cells were rinsed and then mounted on glass slides using Prolong Diamond Antifade reagent Thermo Fisher Scientific).

For HepG2 cells, wells were also fixed for 10 minutes RT with 4% PFA. For immunolabeling experiments, cells were then washed, permeabilized with ice-cold methanol at -20°C for 15 minutes, and then washed again before blocking using 10% goat serum, 1% BSA, and 0.1% Triton X-100 in 1X PBS. To stain for protein targets, Rabbit Anti-LC3B (Cell Signaling Technology, D11 clone, 1:400 dilution), Mouse - Anti-p62/SQSTM1 (Abcam, BSA and azide free, 1:500 dilution), and Chicken Anti - Beta-Actin (Sigma Aldrich, 1:150 dilution) primary antibodies were incubated overnight at 4°C. Secondary antibodies (Goat Anti-Rabbit AlexaFluor 488, Goat Anti-Chicken AlexaFluor 594, and Goat Anti-Mouse AlexaFluor 647, Jackson ImmunoResearch) were incubated for 2 hours at RT. Sample preparation prior to imaging is the same as described above for HEK293A cells.

Freshly fixed samples were blocked before staining for 3 hours with 10% goat serum (Jackson Immuno Research, 005-000-121), 1% bovine serum albumin (Sigma-Aldrich, A9418), and 0.1% Triton-X 100 (Sigma -Aldrich, T8787) in 1X PBS. Primary Antibodies were then incubated overnight at 4°C to label LC3B (1:400, Cell Signaling Technology, 3868), and β-Actin (1:1000, Cell Signaling Technology, 3700), rinsed 3x, and then incubated with secondary antibodies, Goat Anti-Rabbit AlexaFluor 488 (1:2000, Jackson Immuno Research, 111-545-144), and Goat Anti-Mouse AlexaFluor 594 (1:2000, Jackson Immuno Research, 111-585-146). After 3 hours of incubation at RT, nuclei were then labeled by incubating 1µg/mL Hoechst 33342 (Thermo Fisher Scientific, 62249) for 15 minutes at RT, followed by three washes with 1X PBS. Samples were then mounted in Prolong Gold Antifade Reagent (Thermo Fisher Scientific, P36934). For HEK293A cells expressing GFP, samples were mounted in Prolong Diamond Antifade (Thermo Fisher Scientific, P36961). For double labeling experiments, Goat-Anti Rabbit AlexaFluor 647(1:2000, Jackson Immuno Research, 111-605-146) was used to detect LC3B in both the green and the far red channels. To label LC3A and LC3B proteins, a different primary antibody was substituted (1:150, Cell Signaling Technology, 12741S). Samples were normally imaged within a week of staining and kept at 4°C in the dark for preservation.

### Western Blotting

Cells designated for western blotting were grown on 6 well plates or 96 well plates, lifted using cell scrapers, and then lysed with RIPA buffer (Thermo Fisher Scientific, 89900) and protease inhibitor cocktail (Thermo Fisher Scientific, 78430). Western Blot samples were kept frozen at -20°C before analysis.

Protein Extracts were run on 4-20% precast gels, (BioRad, 4569035) at 90V for 2 hrs in Tris-glycine-SDS running buffer, transferred in 25 mM Tris with 192 mM glycine and 20% methanol at 90V for 1 hour. Samples were blocked in 5% non-fat dry milk in TBS-T. Primary antibodies for GAPDH (Cell Signaling Technology, 5174), LC3B (Cell Signaling Technology, 3868), LC3A/B (Cell Signaling Technology, 12741), p62 (Abcam, ab56416), Viniculin (Cell Signaling Technology, 13901), and GFP (Cell Signaling Technology, 2956), were all incubated 1:1000 in 1% Non-fat dry milk in TBS-T. Secondary Antibodies were diluted 1:2000 in 5% BSA in TBS-T solution. Anti-mouse IgG-HRP (Cell Signaling Technology, 7076) and anti-rabbit IgG-HRP (Cell Signaling Technology, 7074) secondary antibodies were diluted 1:2000 in 5% BSA in TBS-T solution. Bands were visualized by chemiluminescence using SuperSignal West Femto Maximum Sensitivity Substrate (Thermo Fisher Scientific, 34094) on a ChemiDoc XRS+ imager (Bio-Rad).

### Preparation of Starved 3D Organoid Samples

For western blot analysis, samples were lysed and stored as described above. For immunofluorescence experiments, organoids were fixed in 4% PFA for 1 hour and then transferred to 30% sucrose solution overnight or until the samples sank to the bottom of the tube. Organoids were then sliced and frozen into 50 µm sections using a microtome. For immunolabeling, organoids were transferred into 1.5 mL tubes. Solution exchanges were achieved by a short spin on a benchtop centrifuge. Samples were mounted using Prolong Gold Antifade. Neurons were identified with Tyrosine Hydroxylase (1:1000 Abcam, ab112), Neuronal marker NeuN (1:500, Abcam, ab177487), and TUJ1 (1:1000, Sigma-Aldrich, MAB1637). Astrocytes were labeled with GFAP (1:500, Sigma-Aldrich, G3893).

### Imaging and Image Analysis

To ensure a quality dynamic range, the gain of each channel was adjusted such that there are no saturated pixels to enable accurate quantitative image analysis^14^. For analysis, we created maximum-intensity Z-projections in ImageJ using a macro and then processed the images simultaneously using CellProfiler^15^. All measurements were made on a per-cell basis, removing cells on the edges of the FOV. The pipelines are available upon request, but we specifically highlight the use of a median filter for nuclei and β-Actin channels to aid in primary and secondary object selection (i.e., nuclei and cell edge segmentation). Additionally, the use of a top hat filter improves the automated detection of relatively small objects (relative to pixel size or resolution) such as LC3 and p62 puncta.

Whole organoid slices for general characterization of neuron population were carried out using widefield microscopy on an Olympus VS120 slide scanner. For 96-well plate formulations, cells were then washed in 1X PBS and then imaged on an Olympus IX-83 widefield microscope and an air immersion 60x, 0.95 NA objective. Mounted slides and live cells were imaged using an Olympus FV3000 confocal microscope and an air immersion 40x, 0.90 NA objective. For LC3 labeled samples, all imaging data was collected on an Olympus FV3000 confocal microscope using a 60x (PLANAPO) oil immersion objective with an NA of 1.42. Cells with different treatment groups were all acquired with the same imaging acquisition parameters. For monoculture cell images, cells were segmented using CellProfiler, with custom steps for nuclei and cell segmentation as needed. For analysis of organoid slides, cell areas specific to neurons (TUJ1+) or astrocytes (GFAP+) were allocated via thresholding of respective channels to create 3D masks, and then applied to the LC3 image for dispersion analysis.

All analysis measurement modules (e.g. MeasureTexture, MeasureAreaShape, etc.) used identical settings across all cell types. For hypoxia experiment analyses, the MeasureObjectIntensityDistribution module was used to quantify the intensity profile of the mitochondria channel from the nuclei to the cell edge. Output data files from CellProfiler pipelines were processed and imported into MATLAB for downstream analysis. To generate dispersion measurements, including linear decomposition, images were directly imported into MATLAB, normalized, and cell regions were identified using counterstains. LC3, mitochondria, and microtubule channel pixels located within cell regions were included in dispersion calculations. Dispersion indices were calculated through the following equations:

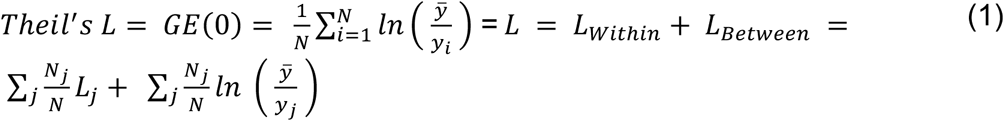

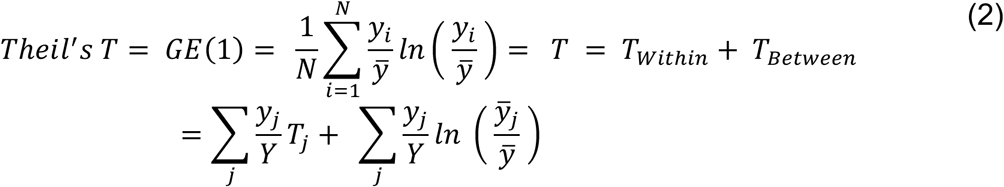

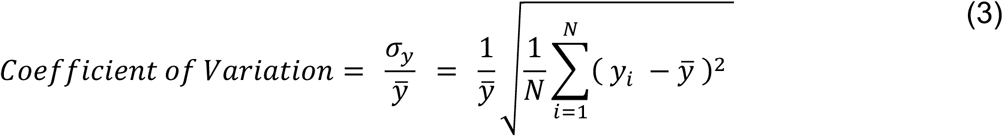

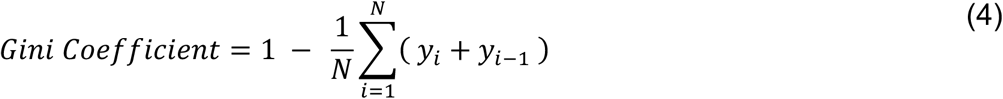

where y_i_ is the intensity of a pixel, y_j_ is the intensity of a pixel within a subgroup, ȳ is the mean intensity of all pixels, ȳ_j_ is the mean intensity of pixels within a subgroup, N is the total number of pixels, σ_2_ is the standard deviation, N_j_ is the total number of pixels in the jth subgroup, Y is the total amount of intensity, Tj is the Theil’s T value calculated considering only subgroup pixels and Lj is the Theil’s L value calculated considering only subgroup pixels.

### Numerical Approximations and Puncta Simulations

For numerical evaluation of dispersion indices, a total intensity was assumed and 99.999% of the total intensity was incrementally distributed from 1 to the total number of pixels (9000), with the remaining percentage distributed evenly among the other pixels. To simulate autophagosome accumulation, puncta were simulated in an approximately 10 µm cell as a 100×100 pixel square cell array with puncta as 2D Gaussians with full width at half maximum (FWHM) equivalent to the range of autophagosomes we normally expect to see in biological data (0.5-1.5µm in diameter). Pixel intensities from puncta were calculated by integration of a 2D Gaussian function. Puncta were assigned random locations in the cell without overlapping centers until no more locations remained. To yield constant total intensity in the simulated image, random noise was overlaid such that the integrated intensity was constant for all puncta diameters and numbers. For assumptions of constant background, the same 2D noise array was added to all simulated images.

### Rank-based performance characterization of dispersion indices

All cell-based measurements were normalized by measurement type and averaged by field of view before performing regression analysis against LC3-II blotting measurements. Regression characteristics for each model were then assigned scores based on 7 different characteristics (fit error, fit variability, fit sensitivity, treatment cross validation, KFold cross validation, R^2^ of fit, and adjusted R^2^) and ranked. Measurements that produced insignificant fits based on a p-value of 0.001 were removed from the downstream rank analysis.

### Statistical Analyses

All plotting and statistical analysis were performed in MATLAB or GraphPad Prism, and statistical tests used unpaired two-tailed t-tests or ANOVA, with Tukey’s or Dunnett’s test for multiple comparisons. P-values less than 0.05 were considered significant. For nonlinear fitting, we performed a 4-parameter (variable slope) fit to estimate EC50 values with 95% confidence intervals for all fitted parameters. The reported slopes are 5x the true slope of the curve due to dose-data transformations that enabled the plotting of box plots on a log_10_ axis and do not affect the quality of the resulting fits.

## Funding

BU Nanotechnology center at Boston University (AM, BT)

National Institutes of Health T32 grant Translational Research in Biomaterials NIH T32EB00635940 (AM, SZ)

National Institutes of Health grant NIH R01CA232056 (MWG)

William Fairfield Warren Distinguished Professorship (MWG)

## Author contributions

Conceptualization: AM, SZ, MWG

Methodology: AM, SZ

Investigation: AM, SZ, AW, BT, MP

Analysis: AM, SZ, DZ

Supervision: MWG

Writing—original draft: AM, SZ, MWG

Writing—review & editing: AM, SZ, AW, BT, MP, DZ, HTN, OS, XH, MWG

## Data and materials availability

All data are available in the main text or the supplementary materials. MATLAB code and CellProfiler pipelines are available at https://github.com/suezhangBU/dispersion_indices.

## Supplementary Figures

**Fig. S1.**
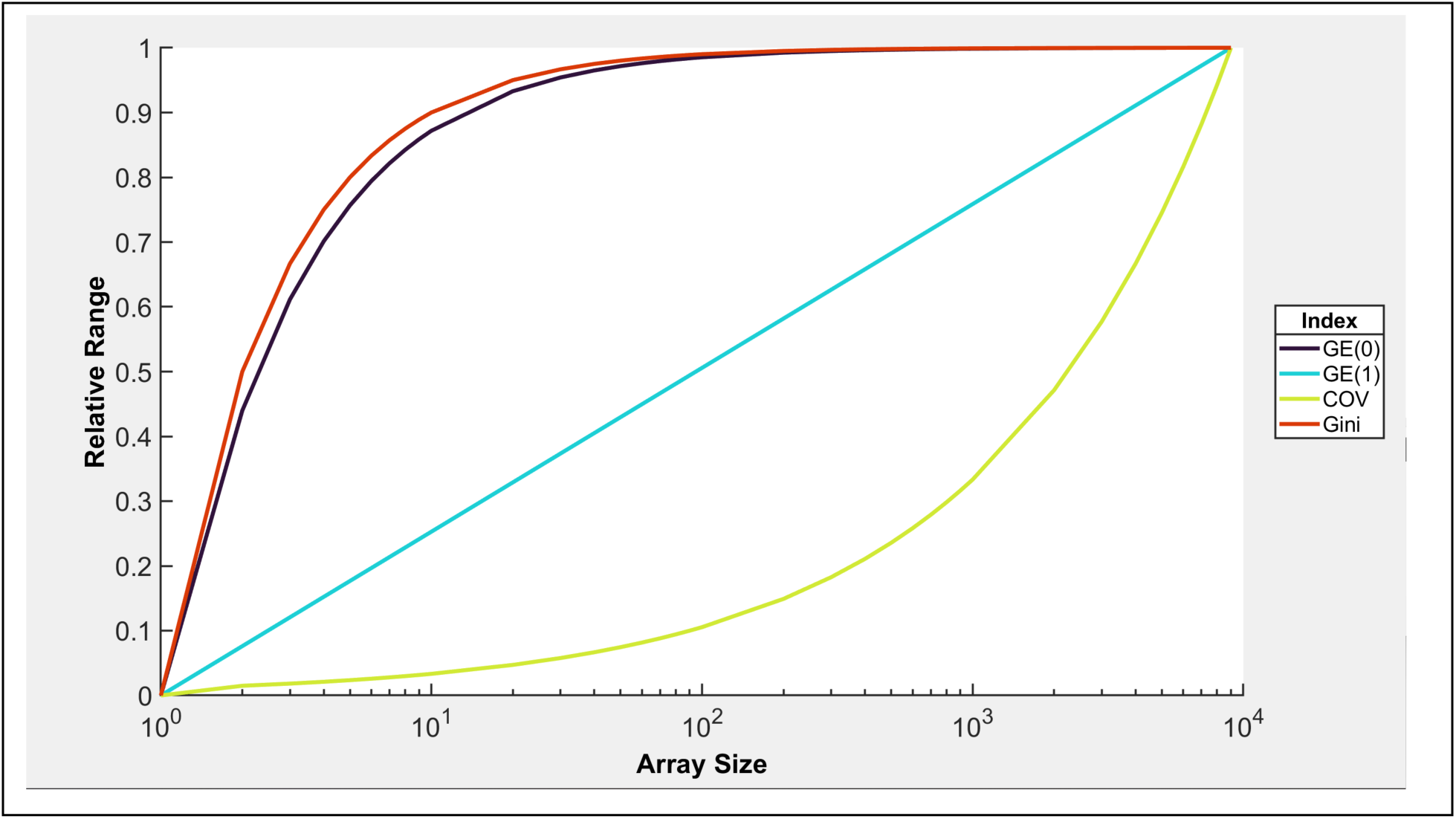
Range of relative values achieved by dispersion indices as a function of array size. Increasing the array size (number of pixels) increasing the possible maximum dispersion index value achieved in cases of high inequality (ie. 99.999% of the pixel intensity is within 1 pixel).

**Fig. S2.**
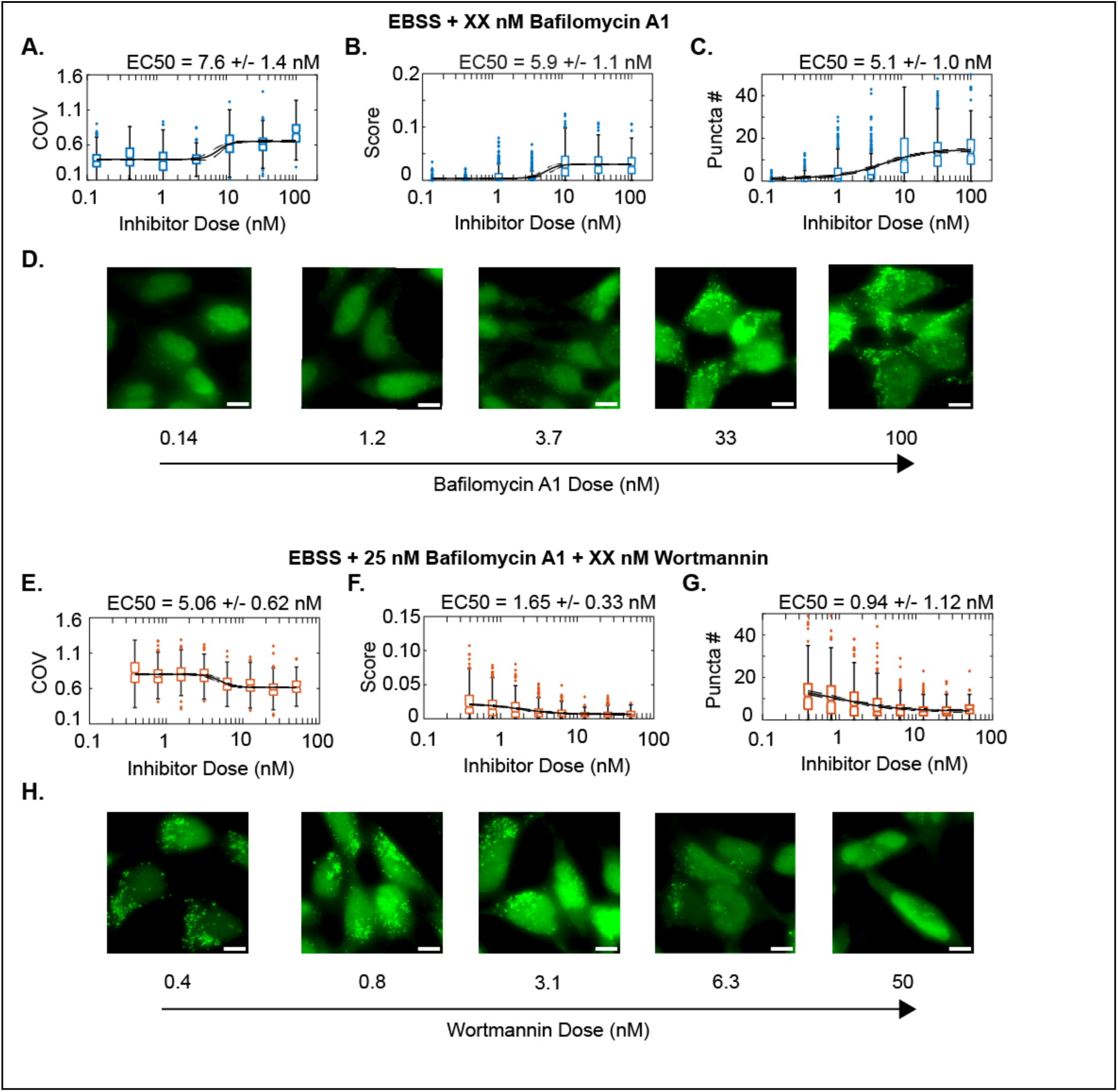
EC50 dose-response generation with HEK293A cells using widefield microscopy. Characterization of bafilomycin A1 response quantified using either the (A) the COV, (B) autophagy score, or (C) automated puncta per cell counting. Representative images for A-C are given in (D). Characterization of wortmannin inhibition of PI3K and autophagy induction given cotreatment with 25 nM bafilomycin A1. Measurements of COV (E), autophagy score (F) and puncta per cell measurements (G) show increasing puncta accumulation with decreasing wortmannin dose. Representative images are presented in (H). Scale bar 5 micron. EC50 values are reported with a 95% CI.

